# Exonuclease III (XthA) enforces *in vivo* DNA cloning of *Escherichia coli* to create cohesive ends

**DOI:** 10.1101/454074

**Authors:** Shingo Nozaki, Hironori Niki

## Abstract

*Escherichia coli* has an ability to assemble DNA fragments with homologous overlapping sequences of 15-40 bp at each end. Several modified protocols have already been reported to improve this simple and useful DNA-cloning technology. However, the molecular mechanism by which *E. coli* accomplishes such cloning is still unknown. In this study, we provide evidence that the *in vivo* cloning of *E. coli* is independent of both RecA and RecET recombinase, but is dependent on XthA, a 3’ to 5’ exonuclease. Here, i*n vivo* cloning of *E. coli* by XthA is referred to as iVEC (*in vivo E. coli* cloning). Next, we show that the iVEC activity is reduced by deletion of the C-terminal domain of DNA polymerase I (PolA). Collectively, these results suggest the following mechanism of iVEC. First, XthA resects the 3′ ends of linear DNA fragments that are introduced into *E. coli* cells, resulting in exposure of the single-stranded 5′ overhangs. Then, the complementary single-stranded DNA ends hybridize each other, and gaps are filled by DNA polymerase I. Elucidation of the iVEC mechanism at the molecular level would further advance the development of *in vivo* DNA-cloning technology. Already we have successfully demonstrated multiple-fragment assembly of up to seven fragments in combination with an effortless transformation procedure using a modified host strain for iVEC.

**Importance:** Cloning of a DNA fragment into a vector is one of the fundamental techniques in recombinant DNA technology. Recently, *in vitro* recombination of DNA fragments effectively joins multiple DNA fragments in place of the canonical method. Interestingly, *E. coli* can take up linear double-stranded vectors, insert DNA fragments and assemble them *in vivo.* The *in vivo* cloning have realized a high level of usability comparable to that by *in vitro* recombination reaction, since now it is only necessary to introduce PCR products into *E. coli* for the *in vivo* cloning. However, the mechanism of *in vivo* cloning is highly controversial. Here we clarified the fundamental mechanism underlying *in vivo* cloning of E. coli and also constructed an *E. coli* strain that was optimized for *in vivo* cloning.

## Introduction

Cloning of a DNA fragment into a vector is one of the fundamental techniques in recombinant DNA technology. As the standard procedure for DNA cloning, a method using restriction enzymes and DNA ligases has long been used. Recently, modified methods of DNA cloning have been widely adopted in place of the canonical method. For example, for the joining of DNA fragments to vectors, an *in vitro* recombination reaction is used. In particular, enzymatic assembly of DNA fragments by using T5 exonuclease, DNA polymerase and DNA ligase effectively joins multiple DNA fragments (1). T5 exonuclease resects the 5’ ends of the terminal overlapping sequences of the DNA fragments to create the 3’ ends of single-stranded DNA overhangs. The complementary single-stranded DNA overhangs are annealed, the gaps are filled, and the nicks are sealed enzymatically. A similar reaction also occurs with the crude cell extract of *Escherichia coli* (2, 3).

In contrast to DNA cloning utilizing *in vitro* recombination, some strains of *E. coli* can take up linear double-stranded vectors, insert DNA fragments and assemble them *in vivo* (4, 5). The ends of these linear DNA fragments need to share 20-50 bp of overlapping sequences with homology. DNA amplification by PCR readily provides this type of linear DNA fragment of interest. Following its introduction, in the early 1990s, this simpler cloning method was not widely used. Recently, however, it has been brought to scientific attention and has been improved with various strains of *E. coli* and several PCR-based protocols (6-12). These improved protocols for *in vivo* cloning have realized a high level of usability comparable to that by *in vitro* recombination reaction, since now it is only necessary to introduce PCR products into *E. coli* for the *in vivo* cloning.

The mechanism of *in vivo* cloning is highly controversial. Initially, the *sbcA23* mutant of the *E. coli* strain JC8679 was used for *in vivo* cloning because the expression of RecE exonuclease and RecT recombinase, here referred to as RecET recombinase, of Rac prophage is activated in this mutation (5, 13). Then, it was thought that a recombination pathway of the prophage was involved in the *in vivo* cloning. However, *E. coli* strains without *sbcA23* mutation, such as DH5α, also have the sufficient ability for *in vivo* cloning (4, 8, 9). Recently, it was suggested that the ability for *in vivo* cloning is not limited to specific mutant strains (10, 11). If *in vivo* cloning is not dependent on host *E. coli* strains, then the DNA substrates may be responsible for the *in vivo* cloning. Klock *et al.* considered that the DNA fragments prepared by PCR have a single-stranded DNA region resulting from incomplete primer extension, and hybridization between complementary single-stranded ends promotes the pathway for *in vivo* cloning (6). On the other hand, Li *et al.* conjectured that 3′ to 5′ exonuclease activity of high-fidelity DNA polymerase creates a single-stranded region at the ends of the linear DNA fragments during PCR (7). Thus, the DNA fragments with single-stranded overhangs produced by PCR seem to be a key for *in vivo* cloning. However, the linear DNA fragments prepared with a restriction enzyme that generates blunt ends are also capable of *in vivo* cloning, indicating that other mechanisms such as a gap repair reaction should be considered (8). In general, the mechanism of *in vivo* cloning remains unclear.

Here we clarified the mechanism underlying the *in vivo* cloning of *E. coli* and also constructed an *E. coli* strain that was optimized for *in vivo* cloning. In addition, we streamlined the procedure of *in vivo* cloning by introducing a newly developed transformation procedure using a single microcentrifuge tube.

## Results

### The iVEC activity in various strains

To identify the principle mechanism underlying the *in vivo* cloning in *E. coli*, here referred to as iVEC, we first confirmed the iVEC activity in various conventionally used strains of *E. coli*. We performed a simple assay of the iVEC activity by transforming the strains with two DNA fragments that carry 20 bp of homologous overlaps at their ends: a *cat* gene encoding chloramphenicol acetyltransferase and the vector plasmid pUC19 (**Fig. 1A**). As a result, transformants resistant to both ampicillin and chloramphenicol appeared in all of the strains tested, although the efficiency of transformation varied depending on the host cells (**Fig. 1B**). MG1655 and JC8679, in particular, had fewer transformants than the other strains. In order to confirm that the *cat* gene was cloned into pUC19, purified plasmids derived from the transformants were analyzed. All of the purified plasmids were larger than the empty vector, pUC19 (**Fig. 1C**). When the plasmids were digested with *Bam*HI, a single band was detected in each lane and the length of the band matched that of the cloned plasmid (**Fig. 1D**). Insertion of DNA into the vector was also confirmed by PCR (**Fig. 1E**).

**Fig. 1.**
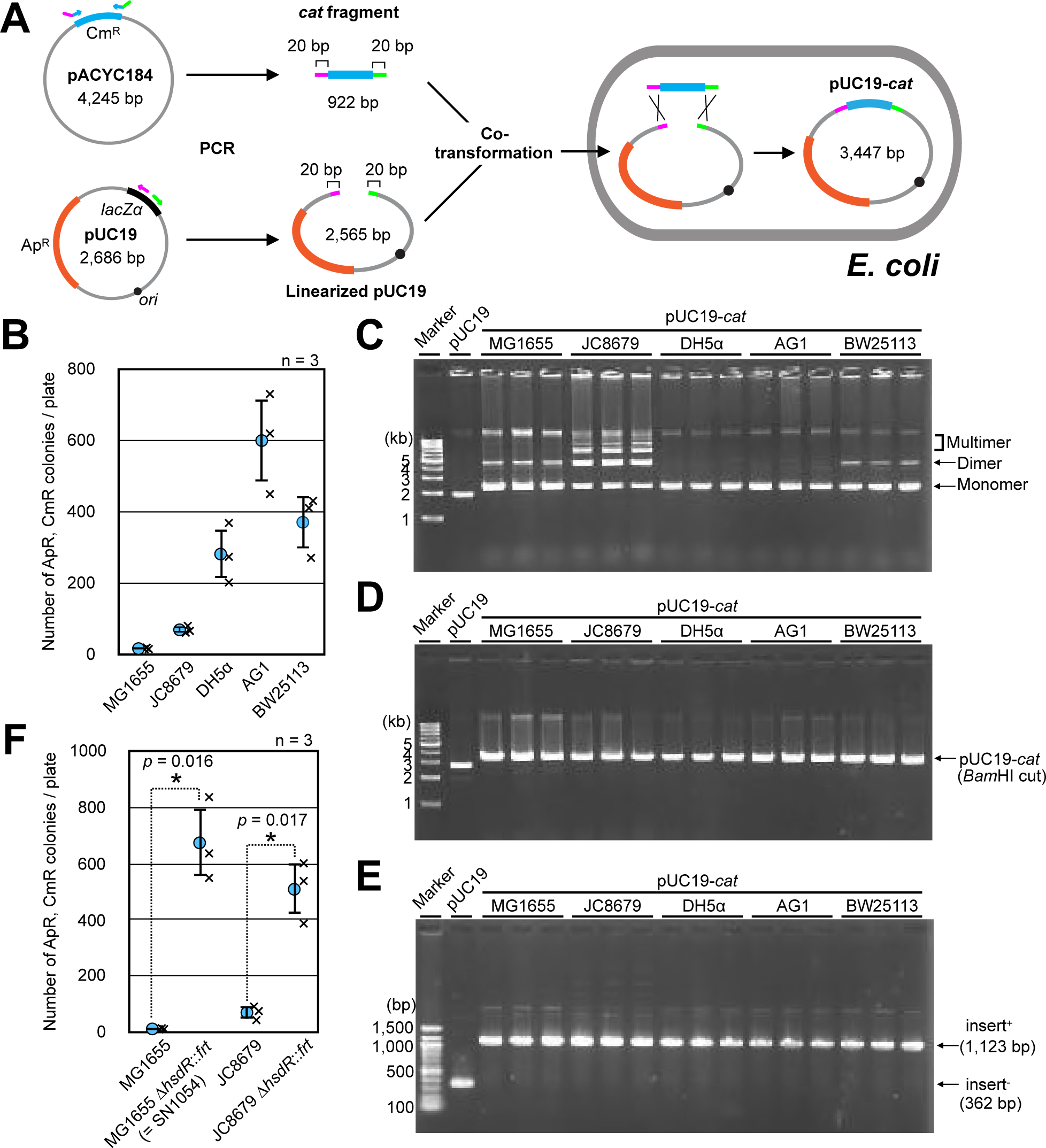
Assays of the iVEC activities. **A.** A scheme of *in vivo* cloning by assembly of two DNA fragments in a cell. DNA fragments containing the *cat* gene and linearized pUC19 DNA have 20 bp homologous overlapping ends (magenta and green). Ampicillin-resistance (Ap^R^) and chloramphenicol-resistance (Cm^R^) genes are shown in orange and light blue, respectively. **B.** The iVEC activities of various strains are shown as the number of colonies resistant to both ampicillin and chloramphenicol. Averages of three independent experiments (crosses) are shown as circles with standard deviations. **C.** Agarose gel electrophoresis of recombinant plasmids that were purified from the indicated strains. Plasmid DNA of pUC19 prepared from DH5α was used as a control. **D.** Agarose gel electrophoresis of the plasmid DNA in (C) after digestion with *Bam*HI. **E.** Confirmation of insert DNA by PCR. The insert sequence was amplified by PCR and the length of PCR products was analyzed by agarose gel electrophoresis. pUC19 without an insert sequence was used as a negative control. **F.** The iVEC activity of strains with Δ*hsdR* mutation. Statistically significant differences are indicated with asterisks (*p value < 0.05 by Welch’s T-test).

Due to the smaller number of positive colonies in MG1655 and JC8679, we noticed that these strains have the wild-type *hsdR* gene. The three other strains, DH5α, AG1 and BW25113, have a mutation in *hsdR.* HsdR is a host specificity restriction enzyme, which degrades DNA containing an unmethylated Hsd recognition sequence (14), and pUC19 DNA contains the recognition sequence. Therefore, we introduced a deletion mutation of the *hsdR* gene into MG1655 and JC8679, resulting in SN1054 and SN1071, respectively. As a result, the numbers of ampicillin‐ and chloramphenicol-resistant colonies after introduction of both the *cat* fragment and linearized pUC19 were significantly increased by the deletion of *hsdR* (**Fig. 1F**). Thus, various *E. coli* strains essentially have the capacity to recombine short homologous sequences at the ends of linear DNAs, permitting the *in vivo* cloning of DNA fragments into linearized vectors.

### *recA* and *recET* are dispensable for the iVEC activity

To elucidate the mechanism of iVEC activity in MG1655, we tested whether recombination proteins such as RecA or RecET were required for the *in vivo* cloning ability. For this purpose, we introduced deletion mutations of the *recA* or *recET* genes into SN1054. We then examined the iVEC activity by transforming these deletion mutants with the *cat* fragment and linearized pUC19. As a result, we found that deletion of *recA* or *recET* had little effect on iVEC activity (**Fig. 2A**), indicating that RecA and RecET are dispensable for *in vivo* cloning.

**Fig. 2.**
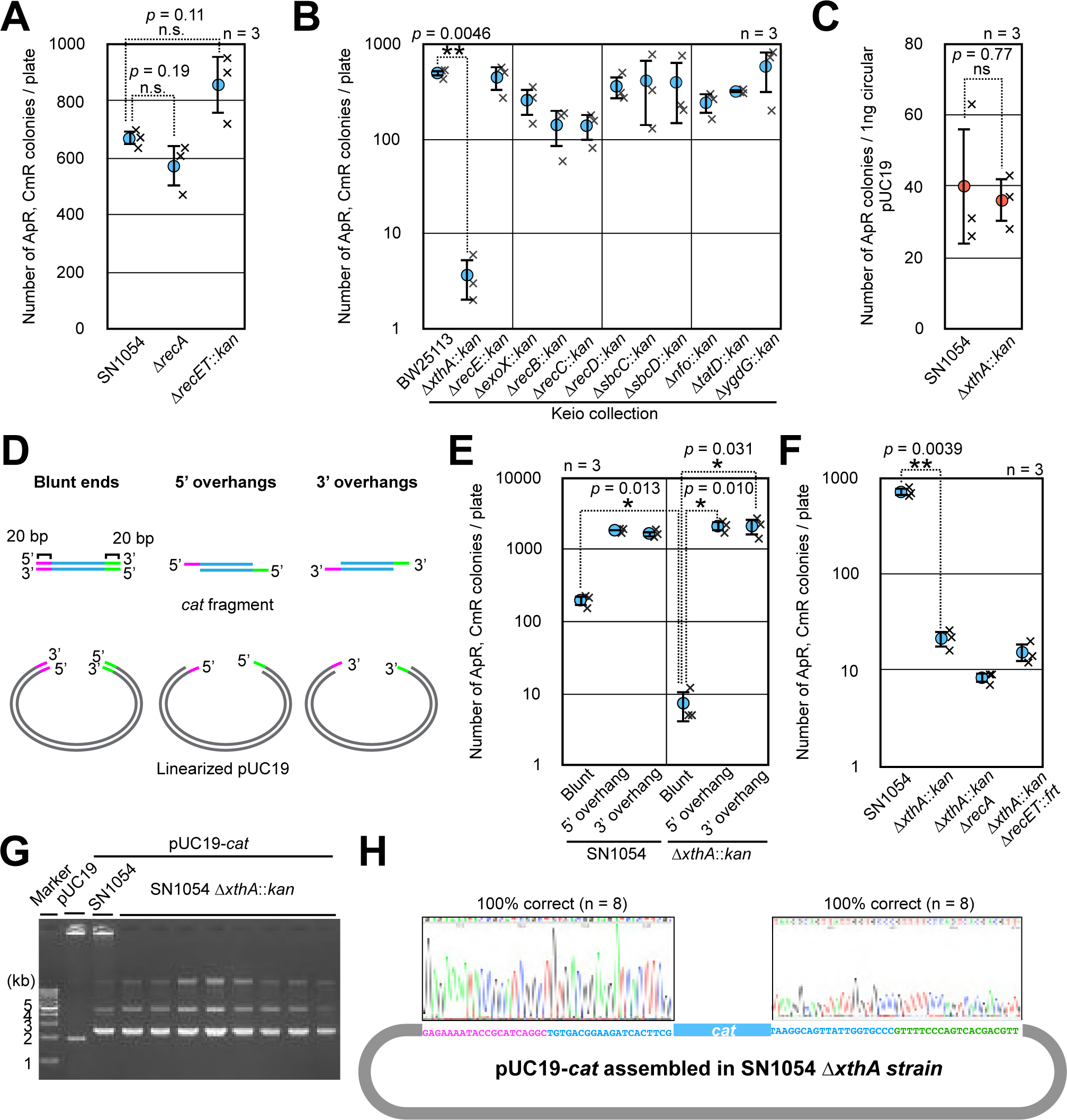
Effect of gene mutations on the iVEC activities. **A.** The iVEC activities of the Δ*recA* and Δ*recET* mutant strains are shown as the numbers of colonies resistant to both ampicillin and chloramphenicol. SN1054 was used as the wild-type strain. Averages of three independent experiments (crosses) are shown as circles with standard deviations. n.s.: not significant (p value > 0.05 by Welch’s T-test). **B.** The iVEC activities of single-gene deletion mutants for various exonucleases in the Keio collection. Asterisks indicate statistically significant differences (**p value = 0.0046 by Welch’s T-test). **C.** Transformation efficiency of the Δ*xthA* strain. One ng of circular pUC19 DNA was used. Averages of three independent experiments (crosses) are shown as circles with standard deviations. ns indicates that the difference is not statistically significant (p value = 0.77 by Welch’s T-test). **D.** A diagram of DNA fragments with blunt ends, 5′ overhangs and 3′ overhangs. *cat* fragments and linearized pUC19 have 20 bp of homologous sequences at ends (magenta and green). **E.** The iVEC activities by using DNA fragments with blunt ends, 5′ overhangs and 3′ overhangs. These DNA fragments were introduced into the SN1054 or Δ*xthA* mutant. Asterisks indicate statistically significant differences (*p value < 0.05 by Welch’s T-test). **F.** The iVEC activities of double gene-deletion mutants: [Δ*xthA* and Δ*recA*] and [Δ*xthA* and Δ*recET*]. Asterisks indicate statistically significant difference (**p value = 0.0039 by Welch’s T-test). **G.** Plasmids assembled in the Δ*xthA* mutant strain were analyzed by agarose gel electrophoresis. pUC19 and pUC19-cat assembled in the *xthA*^+^ strain (SN1054) were used as a control. **H.** Sequencing of the joint region of the plasmids assembled in the Δ*xthA* mutant strain. Eight plasmids of independent single colonies were analyzed.

### *xthA* is required for the iVEC activity

In general, DNA recombination in *E. coli* accompanies conversion of double-stranded DNA to single-stranded DNA by exonuclease. It is reported that *E. coli* has at least seven exonucleases that prefer double-stranded DNA for their substrates as follows: XthA, RecE, ExoX, RecBCD, SbcCD, Nfo and TatD (15). In addition, YgdG is an exonuclease whose preferential substrate is unknown. Next, therefore, we examined the iVEC activity in deletion mutants of these exonucleases. We used the deletion mutants from the Keio collection because BW25113, the parental strain of the Keio collection, has sufficient capacity for iVEC, as shown in Fig. 1B.

We tested each deletion mutant by introducing a DNA fragment containing the *cat* gene and linearized pUC19 vector. As a result, in the Δ*xthA* mutant, the iVEC activity was remarkably decreased to 0.7% of that in the wild-type strain (**Fig. 2B**). The iVEC activity was slightly decreased in the Δ*exoX*, Δ*recB*, Δ*recC*, Δ*nfo* and Δ*tatD* mutants. However, because these defects were several orders of magnitude smaller than that observed in the Δ*xthA* mutant, we focused on XthA in the subsequent experiments.

There was a possibility that deficiency in plasmid maintenance or DNA uptake was the reason for the remarkable reduction of iVEC activity in the Δ*xthA* mutant. Therefore, we examined the level of transformation efficiency of the Δ*xthA* mutant by using circular DNA of the pUC19 plasmid and found that it was almost equivalent to the efficiency of the *xthA*^+^ strain (**Fig. 2C**). This indicates that plasmid maintenance and DNA uptake are normal in the Δ*xthA* strain. Since XthA (exonuclease III) has 3′ to 5′ exonuclease activity (16), we speculated that resection of the DNA ends by this enzyme to produce single-stranded overhangs is crucial for iVEC activity. To confirm this idea, we introduced DNA fragments in which 20 bp of the single-stranded overhangs at the ends were generated in advance, into the Δ*xthA* mutant (**Fig. 2D**). As a result, in the Δ*xthA* mutant, a sufficient number of transformants comparable to the number in the *xthA*^+^ strain were obtained from the DNA fragments with overhangs, whereas DNA fragments with blunt ends yielded few recombinants (**Fig. 2E**). Hybridizing between homologous single-stranded DNA regions of the introduced DNA fragments regardless of 5′ or 3′ overhangs would be essential for recombination of the DNA fragments in the host cell. We concluded that the exonuclease activity of XthA to produce single-stranded overhangs plays a critical role in iVEC activity.

Although XthA is a major factor for the iVEC activity, a small number of recombinant plasmids were still produced in the Δ*xthA* mutant (**Fig. 2F**). The transformants were obtained even when a mutation of Δ*recA* or Δ*recET* was added to the Δ*xthA* mutant. We confirmed that the recombinant plasmids were correctly assembled even in the Δ*xthA* mutant (**Fig. 2G and 2H**). Thus, faint iVEC activity still remained in the Δ*xthA* mutant. These results suggest that there are other minor pathway(s) for iVEC activity, which are independent of XthA, RecA and RecET.

### *polA* affects the iVEC activity

Our results suggested that, following the production of single-stranded DNA segments by XthA, homologous single-stranded DNA segments are hybridized and gaps are produced. We considered that specific DNA polymerases fill the gaps to ligate the hybridized DNA fragments To address which DNA polymerase is involved in gap filling, we examined the effect of defects in DNA polymerases on the iVEC activity. *E. coli* has five DNA polymerases (17). Among them, Pol II, Pol IV and Pol V encoded by *polB*, *dinB* and *umuCD*, respectively, are non-essential for cell growth. Therefore, first, we tested the iVEC activities in the deletion mutants of non-essential DNA polymerases. All of these deletion mutants—i.e., Δ*polB*, Δ*dinB* Δ*umuC* and Δ*umuD*— showed little effect on the iVEC activity (**Fig. 3A**). Thus, these polymerases are not involved in the iVEC activity.

**Fig. 3.**
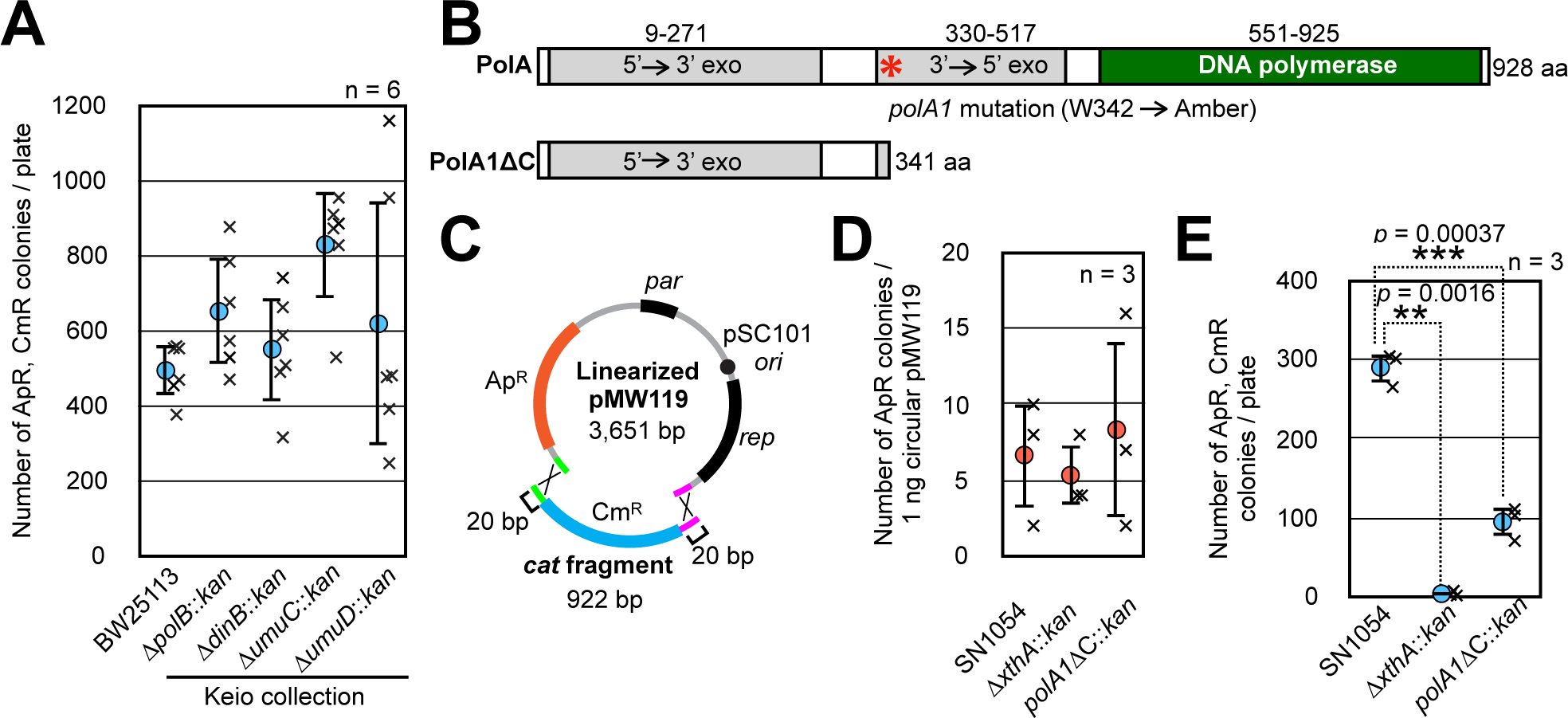
Involvement of DNA polymerases in the iVEC activity. **A.** The iVEC activity of various strains, which are deletion mutants of non-essential polymerases in the Keio collection, are shown as the numbers of colonies resistant to both ampicillin and chloramphenicol. Averages of six independent experiments (crosses) are shown as circles with standard deviations. **B.** A diagram of functional domains in PolA and PolA1 polymerases. An asterisk indicates the point mutation site (W342 to amber) of *polA1* mutation. **C.** Assembly of the *cat* fragment and linearized pMW119 is shown. Each fragment has 20 bp of homologous overlapping sequences shown in green and magenta. **D.** Transformation efficiencies measured by using 1 ng of circular pMW119. Circles indicate averages with standard deviations of three independent experiments (crosses). **E.** The iVEC activity of *polA1*ΔC is shown as the number of colonies resistant to both ampicillin and chloramphenicol after introduction of 0.15 pmol of the *cat* fragment and 0.05 pmol of linearized pMW119 into the indicated strains. Averages of six independent experiments (crosses) are shown as circles with standard deviations. Statistically significant differences compared with the parent strain, SN1054, are indicated with asterisks (**p value = 0.0016 or ***p value = 0.00037 by Welch’s T-test).

Next, we examined the requirement of DNA polymerase I (Pol I) for the iVEC activity. Pol I and Pol III are essential for cell growth. Pol III is a core enzyme of the DNA polymerase III holoenzyme, which is the primary enzyme complex involved in prokaryotic DNA replication. Hence, we considered that it would be difficult to analyze the iVEC activity by using a mutant of pol III. On the other hand, although the *polA* gene encoding Pol I is essential, the full length of this gene is not required for cell viability (18). Only the N-terminal domain encoding 5’ to 3’ exonuclease is sufficient for cell growth (19). Indeed, a *polA1* mutant which expresses only 341 amino acid residues at the N-terminus of PolA by the amber mutation at the amino acid residue 342 is viable (20) (**Fig. 3B**). Accordingly, we constructed a mutant strain carrying the *polA1* mutation, along with deletion of a part of the *polA* gene that encodes the C-terminal 587 amino acid residues including the DNA polymerase domain. The resulting *polA1*ΔC mutant expresses the N-terminal 341 amino acid residues in the manner of the *polA1* mutant. Since the full-length PolA is required for the initiation step of pUC19 replication, we used pMW119 to assay iVEC activity (**Fig. 3C**). The replication origin of pMW119 is derived from pSC101, which does not require the *polA* product for the initiation of its replication (21). The transformation efficiencies of the *polA1*ΔC and the Δ*xthA* mutant with pMW119 were similar to that of a wild-type strain, SN1054 (**Fig. 3D**). We measured the iVEC activity of SN1054 and the Δ*xthA* and *polA1*ΔC mutants by simultaneous introduction of linearized pMW119 and a DNA fragment containing the *cat* gene with a 20 bp overlapping sequence at the ends. High iVEC activity was observed by using pMW119 in the wild-type strain but not in the Δ*xthA* mutant (**Fig. 3E**). Thus, *xthA* played a critical role in the iVEC activity when a pSC101-derivative plasmid vector was used. This result certainly suggests that application of iVEC is not limited to pUC-derivative plasmids. The number of transformants of the *polA1*ΔC mutant decreased to about one third of that of the wild-type strain, and this difference was statistically significant (*p* = 0.00037 by Welch’s T test). In conclusion, the C-terminal domain of PolA was not fully responsible for, but did partly contribute to the iVEC activity.

### Optimization of a host strain for iVEC

Since strains derived from MG1655 had the highest iVEC activity, we attempted to optimize the host strain based on MG1655. Many *E. coli* strains used for DNA manipulation, including DH5α, harbor a mutation in the *endA* gene, which encodes a DNA-specific endonuclease I (22), to improve the quantity of recovered plasmids. Therefore, we introduced a deletion mutation of the *endA* gene into the *E. coli* strain MG1655, along with a deletion mutation of the *hsdR* gene. The number of positive colonies for iVEC increased by two-fold in Δ*endA* cells compared with that of the *endA*^+^ strain (**Fig. 4A**). We examined the transformation efficiency of the Δ*endA* strain with pUC19 plasmid DNA and found that it was increased (**Fig. 4B**). This result indicates that the improvement of the iVEC activity in the Δ*endA* strain was caused by increased transformation efficiency due to the DNA stability during the DNA uptake process.

**Fig. 4.**
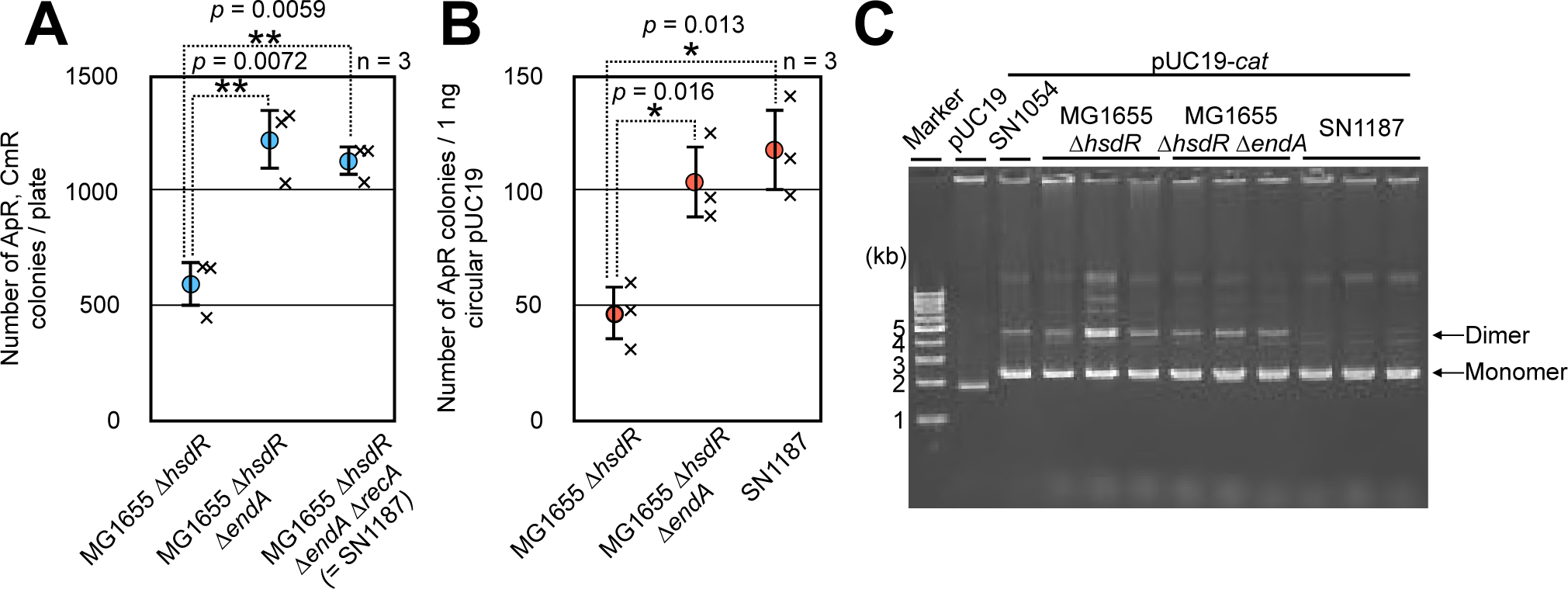
Construction of a strain optimized for iVEC. **A.** Effect of Δ*hsdR* Δ*endA,* and Δ*recA* on the iVEC activities. The iVEC activities are shown as the number of colonies resistant to both ampicillin and chloramphenicol. Averages of three independent experiments (crosses) are shown as circles with standard deviations. Statistically significant differences compared with the MG1655 Δ*hsdR* strain are indicated with askterisks (**p value < 0.01 by Welch’s T-test). **B.** Transformation efficiencies measured by using 1 ng of circular pUC19 in each strain. Averages of three independent experiments (cross) are shown as circles with standard deviations. Statistically significant differences compared with the MG1655 Δ*hsdR* strain are indicated with an asterisk (*p value < 0.05 by Welch’s T-test). **C. A**garose gel electrophoresis of recombinant plasmids (pUC19-*cat*). pUC19 was used as a control vector. The monomer and dimer of the plasmids are indicated as arrows.

In *E. coli*, dimer plasmid DNA is accumulated due to homologous recombination (23). To prevent the dimerization of recombinant plasmids, we introduced a *recA* deletion mutation into a host strain carrying the Δ*hsdR* Δ*endA* strain, resulting in SN1187. Although *recA* deletion mutation often causes lower transformation efficiency due to a reduction in cell viability, the iVEC activity and transformation efficiency of SN1187 were not deteriorated by the deletion mutation of *recA* (**Fig. 4A, B**). Moreover, the amount of dimer was drastically decreased when plasmid DNA was retrieved from SN1187 and analyzed by using agarose gel electrophoresis (**Fig. 4C**).

### Multiple fragment cloning by the host strain SN1187

We further evaluated a new host strain, SN1187, in terms of its capacity for iVEC. First, we examined whether certain lengths of homologous sequences at the ends of DNA fragments were required. We tested DNA fragments with overlapping sequences of 15 bp to 30 bp in length (**Fig. 5A**). In this experiment, the numbers of ampicillin-resistant colonies after introduction of both linearized pUC19 and the *cat* fragment were counted. Approximately 600, 1000, 3200 and 3700 ampicillin-resistant colonies appeared when we used DNA fragments with overlapping sequences of 15 bp, 20 bp, 25 bp and 30 bp at their ends, respectively (**Fig. 5B**). Most of the colonies (99% to 100%) were also resistant to chloramphenicol, indicating that the DNAs were correctly assembled in those colonies (**Fig. 5C**). On the other hand, when only linearized pUC19 was introduced, only 5 ampicillin-resistant transformants appeared (**Fig. 5B**). This result suggests that carryover of a small amount of template vector from PCR yielded few undesirable transformants, despite the fact that *Dpn*I digestion of the template DNA from PCR was not carried out.

**Fig. 5.**
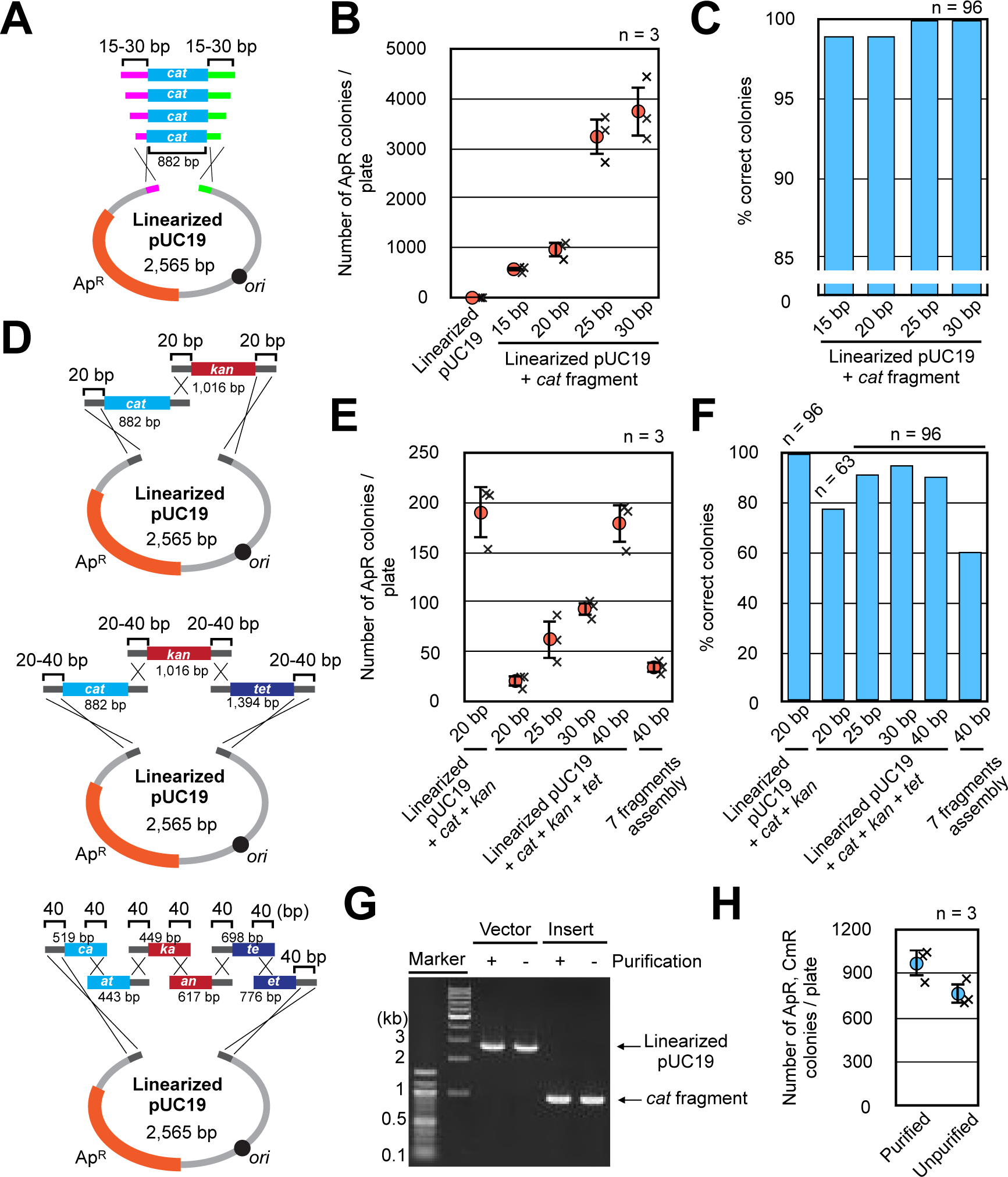
Performance of the iVEC activity by the optimized strain. **A.** A diagram of the assembly of two DNA fragments with varying lengths of overlaps at the ends. **B.** The iVEC activities by using two DNA fragments with varying lengths of overlaps at the ends are shown as the number of colonies resistant to ampicillin. Averages of three independent experiments (crosses) are shown as circles with standard deviations. Introduction of only linearized pUC19 was also carried out as a negative control. **C.** Proportion of colonies which were resistant to chloramphenicol among the 96 ampicillin-resistant colonies in Fig. 5B are shown as the percentage of correct colonies. **D.** A diagram of the assembly of multiple DNA fragments with varying lengths of overlaps at the ends. **E.** The iVEC activities by using multiple DNA fragments with varying lengths of overlaps at the ends are shown as the number of colonies resistant to ampicillin. Averages of three independent experiments (crosses) are shown as circles with standard deviations. **F.** The proportion of colonies that were resistant to antibiotics among the 96 ampicillin-resistant colonies in Fig. 5E are shown as as the percentage of correct colonies correct colonies. Resistance to chloramphenicol and kanamycin was observed for the assembly of three fragments, and resistance to chloramphenicol, kanamycin and tetracycline was observed for the assembly of four and seven fragments of ampicillin-resistant colonies (n = 96 except for assembly of the four DNA fragments with 20 bp overlaps, in which n = 63). **G.** Agarose gel electrophoresis of the PCR products with or without purification, which were used for the assembly of two fragments. **I.** The iVEC activities by using the PCR products with or without purification are shown as the number of colonies resistant to ampicillin. The PCR products were DNA fragments with 20 bp of overlaps at the ends. Averages of three independent experiments (crosses) are shown as circles with standard deviations.

We also examined whether iVEC with SN1187 is available for multi-fragment assembly. First, we introduced three DNA fragments (linearized pUC19 and the DNA fragments including the *cat* or *kan* gene) with 20 bp overlapping sequences at their ends (**Fig. 5D**). Also in this experiment, we selected transformants with only ampicillin resistance, which is a marker of vector DNA, for practical purposes. As a result, about 200 ampicillin-resistant colonies were obtained (**Fig. 5E**). When we examined whether 96 randomly selected, ampicillin-resistant colonies were also resistant to chloramphenicol and kanamycin, we found that all 96 colonies were resistant to chloramphenicol and kanamycin as well as ampicillin (**Fig. 5F**). Next, the assembly of four fragments (linearized pUC19 and the DNA fragments including the *cat, kan,* or *tet* gene) was carried out with 20 to 40 bp of homologous overlapping sequences (**Fig. 5D**). We obtained about 20, 60, 90 and 180 ampicillin-resistant colonies with homologous overlaps of 20, 25, 30 and 40 bp, respectively (**Fig. 5E**). The ratios of colonies resistant to all of ampicillin, chloramphenicol, kanamycin and tetracycline against colonies resistant to ampicillin alone ranged from 80% to 95% (**Fig. 5F**). We also read joint sequences of assembled DNAs to confirm the accuracy of recombination. When 8 plasmids per each construct of two, three and four fragments assembly with 20 bp overlapping sequences were examined, no base change was found within overlapping sequences (**Fig. S1A, S1B, S1C**). Finally, we attempted to perform simultaneous gene assembly of seven fragments. Each of the DNA fragments used for the assembly of four fragments was split and assembled with 40 bp homologous overlaps at its ends (**Fig. 5D**). About 40 colonies resistant to ampicillin were obtained (**Fig. 5E**). Among those ampicillin-resistant colonies, about 60% were also resistant to each of the antibiotics chloramphenicol, kanamycin and tetracycline (**Fig. 5F**). This result indicated that the DNA fragments that included antibiotic-resistance genes separated into 6 fragments were correctly assembled at the same time. We also examined joint sequences of this recombinant plasmid. For this purpose, plasmid DNA from 8 independent colonies was examined. While one plasmid had a 2 bp region of deletion within a joint segment, no base change was found in the other plasmids (**Fig. S1D**). Finally, we demonstrated that purification of the PCR products was not necessary for the iVEC activity. When unpurified PCR products were used directly for iVEC without PCR purification, the number of positive colonies was more than 500 (**Fig. 5G, 5H**). The PCR products can be used easily and relatively quickly without the requirement of any treatments such as column purification, ethanol precipitation or *Dpn*I digestion before transformation.

## Discussion

XthA, also known as exodeoxyribonuclease III, XthA is exodeoxyribonuclease III, exhibits 3’-5’ exonuclease activity. Introducing DNA fragments with cohesive ends into the *E. coli* cells effectively bypasses the requirement of XthA for the iVEC activity (**Fig. 2E**). On the other hand, addition of cohesive ends to insert and vector DNA fragments also strengthens the iVEC activity in wild-type cells (**Fig. 2E**). This is consistent with the previous reports that generation of cohesive ends during PCR is effective for *in vivo* cloning (6, 7). Taken together, these facts indicate that the creation of cohesive ends from the blunt ends of DNA fragments is crucial for the *in vivo* cloning. Therefore, we conclude that XthA exonuclease converts the blunt ends of double-stranded DNA to 5’-protruding ends in the process of the *in vivo* cloning. In consideration of this activity, we propose the following as the most likely mechanism for iVEC as shown in Fig. 6. After the insert and the vector DNA fragments are introduced into the *E. coli* cell, XthA resects the ends of the DNA fragments from the 3’ to 5’ direction, producing 5’ overhanging ends. As the ends of insert and vector DNAs have mutually complementary sequences, the 5’ overhanging ends of the insert and the vector DNA fragments hybridize to each other as cohesive ends. In addition, the gaps are filled by DNA polymerases and the nicks are repaired by DNA ligases. Deletion of the DNA polymerase domain of PolA did not completely abrogate the iVEC activity (**Fig. 3E**). There is a redundant polymerase(s) for the gap filling in iVEC. It is possible that pol II, III, IV or V is involved in the gap filling in the *polA1*ΔC background.

**Fig. 6.**
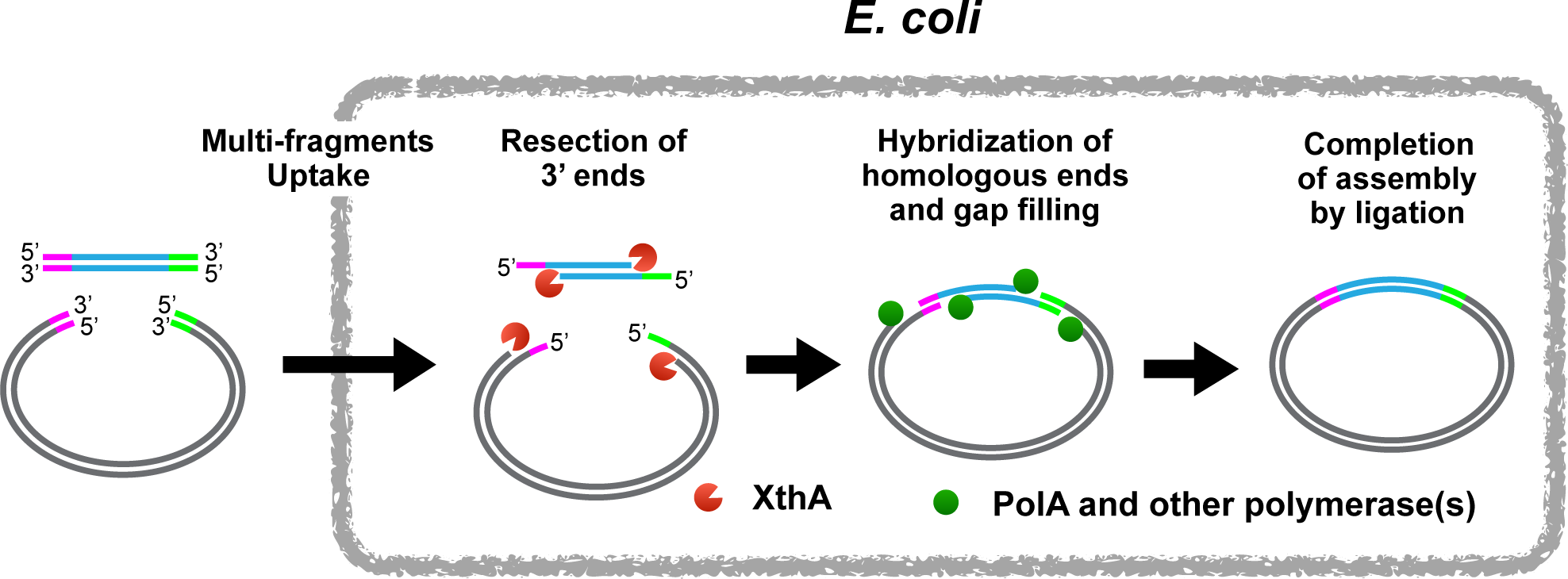
A model for the mechanism of iVEC.

Previously, a strain in which the expression of RecET recombinase was activated by the *sbcA23* mutation was used as a host strain for the *in vivo* cloning (5). Therefore, it was thought that RecET was the recombinase essential for the *in vivo* cloning. While strains without *sbcA23* mutation have been shown to possess the iVEC activity (4, 8, 9), it was not clear whether even a low level expression of RecET was sufficient for iVEC. The present finding that the Δ*recET* mutant exhibited sufficient iVEC activity indicates that RecET is not required for iVEC (**Fig. 2A**). In addition, *E. coli* has other exonucleases in addition to XthA, but their contribution to the iVEC activity is relatively low (**Fig. 2B**). Interestingly, Δ*xthA* cells still maintained slight iVEC activity that was independent of *recA* or *recET* (**Fig. 2F**). This residual activity was not due to PCR-based production of single-stranded overhangs, since it was observed even in the assembly of DNA fragments with blunt ends (**Fig. 2E**). It thus seems likely that some other exonucleases are responsible for the residual iVEC activity in Δ*xthA* cells. XthA would be the dominant exonuclease that preferentially digests double-stranded DNA to produce single-stranded overhangs. Under most conditions, an *E. coli* strain having the exonuclease activity of XthA would be able to assemble DNA fragments with blunt ends that are generated by using a conventional PCR.

Several derivatives of *E. coli* K-12 showed the activity of iVEC, suggesting that no specific mutations are required for the iVEC activity. It seems likely that *E. coli* K-12 originally acquired the iVEC activity, and the iVEC activity was involved in an unknown physiological function in *E. coli*. It is conceivable that XthA would help to repair minor DNA damage, instead of the RecBCD exonuclease. RecBCD produces a 3′ overhang and loads RecA onto the single-stranded DNA, causing an SOS response accompanied by cell division arrest (24). To help avoid such a serious outcome, it is conceivable that XthA could function in a repair pathway of DNA damage.

In our present experiments, we found that the wild-type strain of *E. coli* exhibits iVEC activity, although in general this activity is not high in wild-type strains. To improve the efficiency of iVEC, deletion mutations of *hsdR* and *endA* are introduced. The *hsdR* gene encodes a Type I restriction enzyme, *Eco*KI (25), and EndA is a non-specific DNA endonuclease (22). Both gene disruptions improve the transformation efficiency of the DNA fragments rather than the assembly process. It was expected that enhancement of the expression of *xthA* by using a T5/*lac* promotor would improve the iVEC activity. However, we found that the enhanced expression did not increase the iVEC activity.

We used a modified-TSS method to measure iVEC activity. Cells in overnight culture were used to prepare competent cells for the measurement. Overnight-standing culture allows the entire process to be performed using only a single microcentrifuge tube, from the preparation of competent cells to transformation. In this way, competent cells of many different strains can be easily prepared. However, the transformation of plasmid DNA is not very high: about 10^4^ - 10^5^ CFU/µg pUC19 (**Fig. 4B**). Therefore, by using less than 10-100 pg of template plasmids in PCR products, the background of unwanted vector-only colonies can be significantly reduced. This also means that *Dpn*I treatment after PCR of vector DNA is dispensable in order to reduced transformants by the template plasmid DNA. In fact, we could almost surely obtain the desired colonies despite a lower number of transformants. The number of positive transformants obtained with iVEC using our method and the host strain, SN1187, is comparable or greater than that in previous reports using other methods such as the rubidium chloride method or commercially available competent cells.

Obviously, *E. coli* cells can simultaneously uptake multiple DNA fragments via an unknown mechanism. As a result, assembly of up to seven fragments was possible by using iVEC (**Fig. 5D, 5E, 5F**). In addition, this approach was effective for obtaining recombinant products of less than 10 Kbp in total. To hybridize the cohesive ends of DNA fragments, shorter DNA fragments would be suitable because the opportunity for initial contact between the ends of the DNA fragments increases. At present, our procedure could be utilized for multi-site-directed mutagenesis instead of primer extension mutagenesis. Unexpectedly, single-stranded DNA binding protein (SSB) seemed not to predominantly affect the single-stranded DNA segment that was exposed by XthA. It is conceivable that there is a mechanism to avoid the interference by SSB and promote hybridization between cohesive ends. An improved understanding of the iVEC activity would contribute to the development of iVEC methods in the future.

## Methods

### Medium

L broth (1% Difco tryptone, 0.5% Difco yeast extract, 0.5% NaCl, pH adjusted to 7.0 with 5N NaOH) was used for liquid culture. The agar plate was made of L broth and 1.5% agar. The following antibiotics were used as needed: 50 µg/mL of ampicillin, 10 µg/mL of chloramphenicol, 15 µg/mL of kanamycin and 10 µg/mL of tetracycline.

### Bacterial strains and plasmids

*E. coli* strains and plasmids used in this work are listed in **Table S1 and S2**, respectively. To construct a Δ*hsdR*::*frt* mutant, a chromosomal DNA segment containing Δ*hsdR*::*kan* was amplified from genomic DNA of the Δ*hsdR*::*kan* strain in the Keio collection by PCR using the primer set [hsdR_F and hsdR_R] (26). The amplified DNA fragments were introduced into the parent strains with pKD46 as described by Datsenko and Wanner (27). The Δ*xthA*::*kan*, Δ*recET*::*kan* and *polA1*ΔC::*kan* strains were constructed in a similar manner using the primer sets and templates [xthA_F, xthA_R and chromosome of Keio Δ*xthA*::*kan*], [recET_F, recET_R and pKD4] and [polAdelC_F, polAdelC_R and pKD4], respectively. The *kan* cassette was removed by pCP20, if needed (27). To construct a Δ*recA* strain, a plasmid DNA of pKH5002SB was amplified by using the primer set [pKH_F and pKH_R]. Upstream and downstream chromosomal segments of the *recA* gene were amplified from MG1655 genomic DNA by using the primer sets [recAup_F and recAup_R] and [recAdown_F and recAdown_R]. We obtained a 1.8 kb upstream chromosomal segment and a 2 kb downstream chromosomal segment of recA, respectively. Both the recAup_F primer and the recAdown_R primer have an additional 20 bp complementary sequence complementary to primers pKH_R and pKH_F, respectively. In addition, 40 bp of a sequence within the primers recAup_R and recAdown_F are complementary to each other. Amplified DNA fragments of pKH5002SB, the upstream and the downstream regions of chromosomal segment of *recA* were introduced into a Δ*rnhA*::*kan* strain to generate pKH5002SBΔ*recA* (**Fig. S2A**). Using this plasmid, the *recA* gene was deleted with two successive homologous recombinations as described previously (28) (**Fig. S2B**). The Δ*hsdR* and Δ*endA* strains were constructed by using the same method with the primer sets [hsdRup_F, hsdRup_R, hsdRdown_F and hsdRdown_R] and [endAup_F, endAup_R, endAdown_F and endAdown_R], respectively.

### Preparation of PCR products for transformation

We used KOD plus Neo (TOYOBO) for PCR. The thermal cycler program was as follows: 94 °C for 2 min, followed by 30 cycles of [98 °C for 10 sec, 58 °C for 10 sec, and 68 °C for 30 sec/kb], and a final extension of 68 °C for 5 min. Oligonucleotide primers used for PCR are listed in **Table S3** and **S4**. The final concentration of the template DNA in each reaction mixture was adjusted to 1 pg/µL, e.g., 50 pg in a 50 µL reaction. The *cat* (chloramphenicol-resistance) and *tet* (tetracycline-resistance) genes were amplified from pACYC184 DNA, and the *kan* (kanamycin-resistance) gene was amplified from pACYC177 DNA. All PCR products were purified using a Wizard SV PCR Clean-Up System (Promega). Digestion of template DNA by *Dpn*I was not necessary after PCR.

### Preparation of DNA fragments with blunt ends, 5′ overhangs or 3′ overhangs

DNA fragments with blunt ends, 5′ overhangs or 3′ overhangs were prepared as follows. To isolate single-stranded strands, we used a Long ssDNA Preparation kit (BioDynamics Laboratory, Tokyo). Plasmids used for the isolation of ssDNAs are listed in **Table S2**. Each pair of the top and the bottom single-stranded DNA fragments for blunt ends, 5′ overhangs or 3′ overhangs was mixed and incubated at 99 °C for 5 minutes and annealed at 65 °C for 30 minutes to generate double-stranded DNA.

### Transformation

To introduce DNA fragments into *E. coli* cells, we used the TSS method with modification (29). A small number of cells in a colony on an agar plate was picked up using a sterilized toothpick and suspended in a 1.5 mL microcentrifuge tube containing 1 mL of L broth. The tube lid was closed. The tube was standing in an incubator at 37 °C for 20 hours without shaking. After standing incubation for 20 hours, the OD_600_ of the culture reached approximately 1.4 and the number of cells in the tube was about 4 x 10^8^ CFU/mL. The tube was chilled on ice for 10 minutes and centrifuged at 5,000 g for 1 minute at 4 °C to spin down the cells. The supernatant was removed, and the cell pellet was dissolved in 100 µL of ice-cold TSS solution (50% L broth, 40% 2xTSS solution and 10% DMSO) mixed with DNA. The composition of 2xTSS solution was [20% (w/v) PEG8000, 100 mM MgSO_4_ and 20% (v/v) glycerol in L broth]. For DNA cloning, 0.05 pmol of linearized vector and 0.15 pmol of each insert DNA fragment were used. After gentle mixing, the solution was immediately frozen in liquid nitrogen for 1 minute. Frozen tubes were transferred to an ice bath. After 10 minutes of incubation on ice, the tubes were briefly vortexed to mix their contents and incubated on ice for an additional 10 minutes. Then, 1 mL of L broth was added, and the contents of the tube were mixed by inversion and incubated at 37 °C for 45 minutes. After incubation, the cells were centrifuged and the supernatant was roughly discarded. The cell pellet was dissolved in the remaining supernatant and the cell suspension was spread on an L agar plate containing appropriate antibiotics. Finally, the plates were incubated at 37 °C for 16 hours and the number of colonies was counted. To examine transformation efficiency, 1 ng of the indicated circular plasmids was used.

### Assay of the iVEC activity

DNA fragment containing an antibiotic-resistance gene and linearized pUC19 with 20 bp homologous overlapping ends were amplified by PCR and introduced into *E. coli* cells by modified-TSS method as described above (**Fig. 1A**). In a standard assay of the iVEC activity, 0.15 pmol of *cat* fragment and 0.05 pmol of linearized pUC19 were used for transformation of indicated strains. We counted number of colonies resistant to both ampicillin and chloramphenicol after simultaneous introduction of *cat* fragment and linearized pUC19 into indicated strains.

## Acknowledgements

We thank Dr. Katsuhiro Hanada for the critical suggestions on *in vivo* cloning. We thank NBRP *E. coli* for providing *E. coli* strains and plasmids. This work was supported by a JSPS KAKENHI Grant (no. 8K19193).

## Figure legends

**Fig. S1 Sequencing of the joint region of the assembled plasmids in SN1187.** Eight plasmids of each construct from independent single colonies were analyzed. Primers used for the sequencing reaction and the percentages of correct sequences are shown.

**A.** Joint sequence of plasmids constructed by the assembly of two fragments with 20 bp homologous overlaps.

**B.**Joint sequence of plasmids constructed by the assembly of three fragments with 20 bp homologous overlaps.

**C.**Joint sequence of plasmids constructed by the assembly of four fragments with 20 bp homologous overlaps.

**D.**Joint sequence of plasmids constructed by the assembly of seven fragments with 40 bp homologous overlaps. A 2 bp region of deletions observed in one of the plasmids is indicated with arrows.

**Fig. S2 Construction of deletion mutant by two successive homologous recombinations.**

**A.** Construction of the targeting vector. Linearized pKH5002SB and the upstream and downstream sequences of the target gene were prepared by PCR and assembled in the Δ*rnhA* strain. pKH5002SB could be replicated only in RnaseH-deficient strains, due to deletion of the HaeIII fragment in its replication origin.

**B.** Deletion of the target gene by two successive homologous recombinations. Since pKH5002SB can be replicated only in RnaseH-deficient strains, the plasmid sequence is not maintained as a plasmid but is maintained in a chromosomally integrated state when the plasmid is introduced into the *rnhA*^+^ strains. Cells in which the plasmid sequence is integrated into chromosome are selected by ampicillin. *E. coli* cells harboring the *sacB* gene are not viable on an agar plate containing sucrose, and therefore cells in which the plasmid sequence is dropped out are selected on the sucrose plate.

## References

1. Gibson DG, Young L, Chuang R-Y, Venter JC, Hutchison CA, Smith HO. 2009. Enzymatic assembly of DNA molecules up to several hundred kilobases. Nat Methods 6:343–345.

2. Zhang Y, Werling U, Edelmann W. 2012. SLiCE: a novel bacterial cell extract-based DNA cloning method. Nucleic Acids Res 40:e55.

3. Motohashi K. 2015. A simple and efficient seamless DNA cloning method using SLiCE from Escherichia coli laboratory strains and its application to SLiP site-directed mutagenesis. BMC Biotechnol 15:47.

4. Bubeck P, Winkler M, Bautsch W. 1993. Rapid cloning by homologous recombination in vivo. Nucleic Acids Res 21:3601–3602.

5. Oliner JD, Kinzler KW, Vogelstein B. 1993. In vivo cloning of PCR products in E. coli. Nucleic Acids Res 21:5192–5197.

6. Klock HE, Koesema EJ, Knuth MW, Lesley SA. 2008. Combining the polymerase incomplete primer extension method for cloning and mutagenesis with microscreening to accelerate structural genomics efforts. Proteins 71:982–994.

7. Li C, Wen A, Shen B, Lu J, Huang Y, Chang Y. 2011. FastCloning: a highly simplified, purification-free, sequence‐ and ligation-independent PCR cloning method. BMC Biotechnol 11:92.

8. Jacobus AP, Gross J. 2015. Optimal cloning of PCR fragments by homologous recombination in Escherichia coli. PLoS ONE 10:e0119221.

9. Kostylev M, Otwell AE, Richardson RE, Suzuki Y. 2015. Cloning Should Be Simple: Escherichia coli DH5α-Mediated Assembly of Multiple DNA Fragments with Short End Homologies. PLoS ONE 10:e0137466.

10. Beyer HM, Gonschorek P, Samodelov SL, Meier M, Weber W, Zurbriggen MD. 2015. AQUA Cloning: A Versatile and Simple Enzyme-Free Cloning Approach. PLoS ONE 10:e0137652–20.

11. García-Nafría J, Watson JF, Greger IH. 2016. IVA cloning: A single-tube universal cloning system exploiting bacterial In Vivo Assembly. Sci Rep 6:27459.

12. Huang F, Spangler JR, Huang AY. 2017. In vivo cloning of up to 16 kb plasmids in E. coli is as simple as PCR. PLoS ONE 12:e0183974.

13. Gillen JR, Willis DK, Clark AJ. 1981. Genetic analysis of the RecE pathway of genetic recombination in Escherichia coli K-12. J Bacteriol 145:521–532.

14. Sain B, Murray NE. 1980. The hsd (host specificity) genes of E. coli K 12. Mol Gen Genet 180:35–46.

15. Lovett ST. 2011. The DNA Exonucleases of Escherichia coli. EcoSal Plus 4:1–45.

16. Demple B, Johnson A, Fung D. 1986. Exonuclease III and endonuclease IV remove 3’ blocks from DNA synthesis primers in H2O2-damaged Escherichia coli. Proc Nat Acad Sci 83:7731–7735.

17. Fijalkowska IJ, Schaaper RM, Jonczyk P. 2012. DNA replication fidelity in Escherichia coli: a multi-DNA polymerase affair. FEMS Microbiol Rev 36:1105–1121.

18. De Lucia P, Cairns J. 1969. Isolation of an E. coli strain with a mutation affecting DNA polymerase. Nature 224:1164–1166.

19. Kornberg A, Baker TA. 1992. DNA replication.

20. Joyce CM, Kelley WS, Grindley ND. 1982. Nucleotide sequence of the Escherichia coli polA gene and primary structure of DNA polymerase I. J Biol Chem 257:1958–1964.

21. Timmis K, Cabello F, Cohen SN. 1974. Utilization of two distinct modes of replication by a hybrid plasmid constructed in vitro from separate replicons. Proc Nat Acad Sci 71:4556–4560.

22. Lehman IR, Roussos GG, Pratt EA. 1962. The deoxyribonucleases of Escherichia coli. II. Purification and properties of a ribonucleic acid-inhibitable endonuclease. J Biol Chem 237:819–828.

23. Summers DK, Beton CW, Withers HL. 1993. Multicopy plasmid instability: the dimer catastrophe hypothesis. Mol Microbiol 8:1031–1038.

24. Churchill JJ, Anderson DG, Kowalczykowski SC. 1999. The RecBC enzyme loads RecA protein onto ssDNA asymmetrically and independently of chi, resulting in constitutive recombination activation. Genes Dev 13:901–911.

25. Murray NE. 2000. Type I restriction systems: sophisticated molecular machines (a legacy of Bertani and Weigle). Microbiol Mol Biol Rev 64:412–434.

26. Baba T, Ara T, Hasegawa M, Takai Y, Okumura Y, Baba M, Datsenko KA, Tomita M, Wanner BL, Mori H. 2006. Construction of Escherichia coli K-12 in-frame, single gene knockout mutants: the Keio collection. Mol Syst Biol 2:2006.0008.

27. Datsenko KA, Wanner BL. 2000. One-step inactivation of chromosomal genes in Escherichia coli K-12 using PCR products. Proc Nat Acad Sci 97:6640–6645.

28. Kitagawa R, Ozaki T, Moriya S, Ogawa T. 1998. Negative control of replication initiation by a novel chromosomal locus exhibiting exceptional affinity for Escherichia coli DnaA protein. Genes Dev.

29. Chung CT, Niemela SL, Miller RH. 1989. One-step preparation of competent Escherichia coli: transformation and storage of bacterial cells in the same solution. Proc Nat Acad Sci 86:2172–2175.

